# A comparison between single cell RNA sequencing and single molecule RNA FISH for rare cell analysis

**DOI:** 10.1101/138289

**Authors:** Eduardo Torre, Hannah Dueck, Sydney Shaffer, Janko Gospocic, Rohit Gupte, Roberto Bonasio, Junhyong Kim, John Murray, Arjun Raj

## Abstract

The development of single cell RNA sequencing technologies has emerged as a powerful means of profiling the transcriptional behavior of single cells, leveraging the breadth of sequencing measurements to make inferences about cell type. However, there is still little understanding of how well these methods perform at measuring single cell variability for small sets of genes and what “transcriptome coverage” (e.g. genes detected per cell) is needed for accurate measurements. Here, we use single molecule RNA FISH measurements of 26 genes in thousands of melanoma cells to provide an independent reference dataset to assess the performance of the DropSeq and Fluidigm single cell RNA sequencing platforms. We quantified the Gini coefficient, a measure of rare-cell expression variability, and find that the correspondence between RNA FISH and single cell RNA sequencing for Gini, unlike for mean, increases markedly with per-cell library complexity up to a threshold of ∼2000 genes detected. A similar complexity threshold also allows for robust assignment of multi-genic cell states such as cell cycle phase. Our results provide guidelines for selecting sequencing depth and complexity thresholds for single cell RNA sequencing. More generally, our results suggest that if the number of genes whose expression levels are required to answer any given biological question is small, then greater transcriptome complexity per cell is likely more important than obtaining very large numbers of cells.

## Introduction

Single cell biology has exploded in recent years to touch many areas of biomedical research, stemming from a growing appreciation that individual cells may deviate from the population average [1–3]. Such deviations may arise from the mixture of different cell “types” or cell states within a given tissue, or may arise from probabilistic behavior within a single cell type. Two classes of expression profiling have emerged in single cell analysis. One is single cell RNA sequencing, which gives enormous breadth by measuring the entire cellular transcriptome [4,5]. Another is in situ hybridization approaches such as single molecule RNA FISH [6–8], which yields highly accurate RNA counts and localization data, but typically just for a more limited set of genes.

Single cell RNA sequencing has, in particular, emerged as a potent tool for classifying cells into different “types”, including rare cell types [9], owing to the fact that individual cell types vary in the expression of a large number of genes. This high dimensional redundancy allows the classification to tolerate a large degree of technical inaccuracy: for instance, many genes can be incorrectly read out as a zero [10], but the remaining genes will still give enough data to correctly classify the cell type. As such, reports have demonstrated that even relatively shallow coverage for most cells is sufficient for this application [11].

There are situations, however, in which cells may not necessarily separate into cell types with large numbers of expression differences, but instead may vary in the expression of just a few genes. Examples include rare-cell biology in otherwise clonal cell types, distinguishing more narrowly defined cell states such as cell cycle phase or stress responses, and discriminating very similar cell subtypes that differ only in the expression of a few genes. In those situations, the technical issues surrounding single cell RNA sequencing may have more consequences, but most previous studies [12–15] (with the notable exception of Grün et al. [16]) lacked the gold-standard reference needed to know what degree of data quality, such as degree of transcriptome coverage, is required in order to make robust biological inferences.

Here, we compare high throughput single cell RNA sequencing to RNA FISH to make principled estimates of the quality of single cell RNA sequencing required to recapitulate important features of gene expression like rare-cell expression. Using a melanoma cell line in which we have measured 26 genes with RNA FISH in many thousands of cells as a model, we performed single cell RNA sequencing using both DropSeq and Fluidigm’s C1 mRNA Seq HT IFC platform. We found that while single cell RNA sequencing methods agree with RNA FISH for detecting average levels of expression, other aspects of single cell expression measurements (such as rare cell analysis) required subjecting single cell data to stringent quality control metrics for transcriptome coverage. We show that these metrics are also important for analyses of cellular states that involve several genes, such as cell cycle.

## Results

First, we describe the three methods in our comparison. We performed RNA FISH on WM989-A6, a clonal isolate of the melanoma line WM989. This cell line exhibits drug resistance to vemurafenib at a frequency of around 1:2000 to 1:5000 cells. It also exhibits “jackpot” expression (i.e., rare cells with high levels of expression with the rest having very little) for a number of genes strongly associated with resistance to Vemurafenib, including EGFR, AXL, WNT5A, and NGFR, at frequencies between ranging from one in 50-500 cells (Shaffer et al. Nature, in press). We performed RNA FISH for 17 resistance markers and 9 housekeeping/ubiquitously expressed genes in 7,000-88,000 cells, depending on the gene (Fig. 1A).

**Figure 1.**
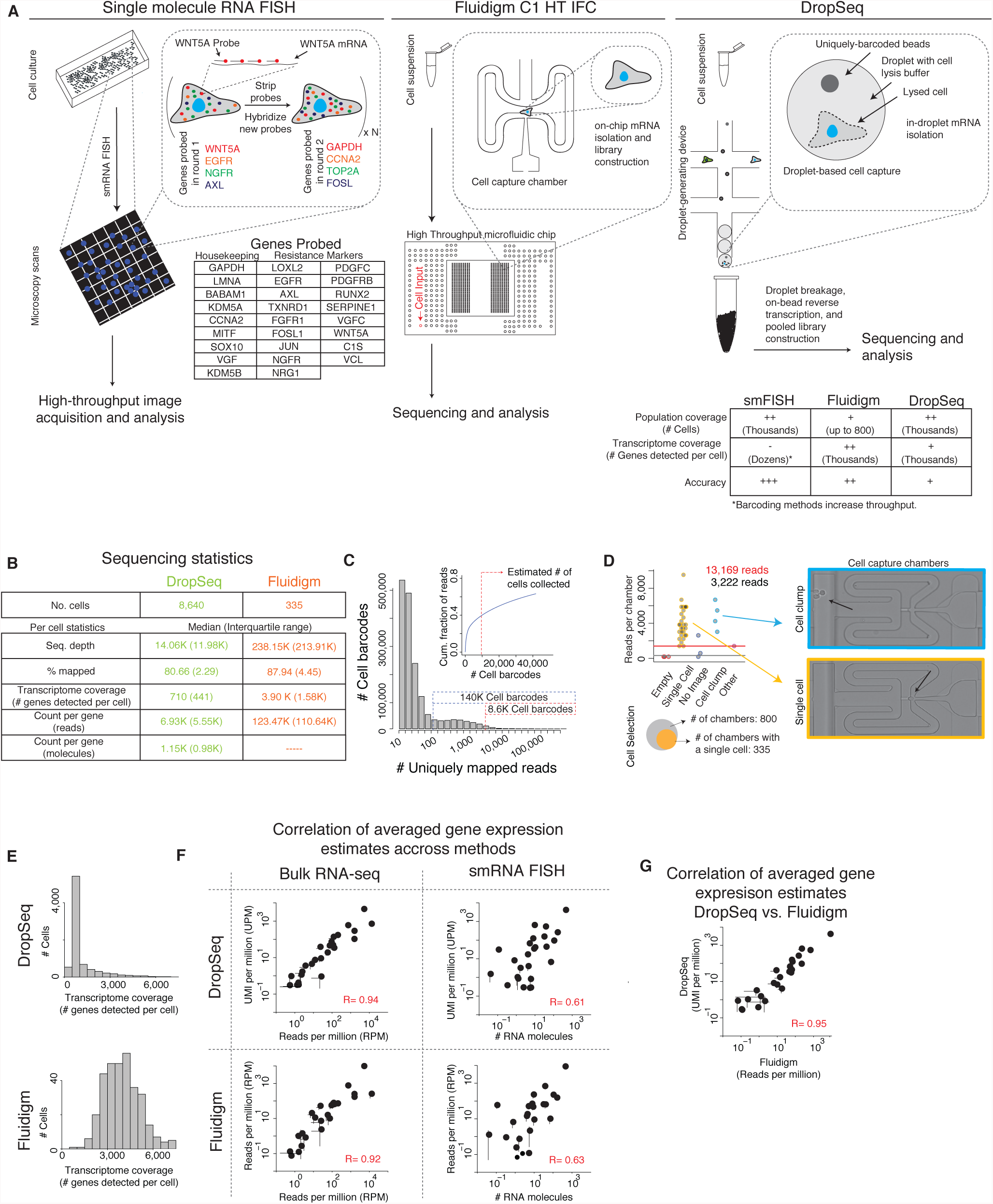
Gene expression estimates correlate between platforms despite differences probing scheme and transcriptome coverage. (A) Schematic of how single molecule RNA FISH (left), Fluidigm’s C1 HT IFC (center), and DropSeq (right) estimate gene expression from individual cells. (B) Sequencing statistics for libraries built with DropSeq (green, n = 1 biological replicate) and Fluidigm (orange, n = 1 biological replicate). (C) Distribution of reads across all barcodes sequenced in DropSeq. (D) Distribution of reads across capture chambers in Fluidigm’s platform. (E) Distribution of transcriptome coverage (# genes detected per cell) for DropSeq (top) and Fluidigm (bottom). (F) Correlation of averaged gene expression estimates across Fluidigm, DropSeq, and bulk RNA-seq. (G) Correlation of averaged gene expression estimates between DropSeq and Fluidigm. Error bars in (F,G) represent two times the standard error of the mean (SEM).

We performed DropSeq on WM989-A6-G3, a clonal isolate of WM989, as per Macosko et al. [17]. DropSeq involves running cells through a microfluidic droplet maker that randomly produces droplets containing individual cells along with a barcoded RNA capture bead. Cells lyse within the droplet, and the RNA adheres to the oligonucleotides on the bead. These bead oligonucleotides contain two codes: one unique to the bead for cell identification, and one unique to each individual oligonucleotide to serve as a unique molecular identifier (UMI). The droplets in the emulsion are then broken and the beads pooled before reverse transcription occurs using the bead oligonucleotides as templates. We then pool the resulting cDNA for library construction and subsequent sequencing (Fig. 1B). We performed DropSeq on human/mouse cell mixtures to confirm that the method was able to robustly separate individual cells in our hands (Supp Fig. 1A). Based on the number of beads used for library construction, we expected to obtain roughly 8600 melanoma transcriptomes and the top 8600 barcodes had at least 5000 uniquely mapped reads per barcode. We found the large number of remaining barcodes formed a bimodal distribution, with ∼140,000 barcodes between 100-5000 uniquely mapped reads, and ∼1.4 million barcodes with <100 uniquely mapped reads. These may correspond to background from free beads either from cells lysed in droplets without a bead or transferring from one bead to another, and from sequencing or PCR errors (Fig. 1C). For the remaining analysis we included only the top 8600 barcodes, with a median depth of 6,938 uniquely mapped reads per cell and an interquartile range (IQR) of 5,553 reads.

We also used Fluidigm’s C1 mRNA Seq HT chip on WM989-A6-G3 cells to generate single cell RNA sequencing data. This method uses microfluidics to load individual cells into 800 separate chambers, lyses the cells, and then performs SMARTer v3 chemistry for RNA amplification. It barcodes and pools the 800 wells into 20 groups of 40, and we generated sequencing libraries for each of the 20 groups separately (no UMIs are included as part of the protocol). We obtained brightfield images for most chambers after cell loading and excluded chambers that were either empty, had clumps of cells, or contained debris in addition to cells (Fig. 1D), leaving 340 potential cells. Of those, we included in our analysis only chambers with more reads than empty chambers without cells, leaving us with data from 335 of the 800 possible chambers. These were sequenced to a median depth of ∼123,000 uniquely mapped reads per cell with an IQR of 110,642 reads.

We then analyzed the quality of the data from both the DropSeq and Fluidigm platforms. We found that as we increased our sequencing depth the number of new genes detected begins to plateau, which suggests that if were we to sequence our libraries more deeply, the majority of the reads would be from amplicons already sequenced (downsampling analysis in Supp. Fig. 1B). As expected, we found generally high levels of expression for melanocyte specific genes such as Sox10, UCN2, MLANA, and DCT but negligible levels of markers for pancreas, heart and spleen (Supp. Fig 1C). Thus, we concluded that our single cell RNA sequencing libraries provided transcriptome measurements consistent with the melanocyte cell line (see also comparison to bulk RNA sequencing data; Fig. 1F).

While these bulk metrics provided some assurance in the quality of our DropSeq data, at the single cell level, we noted wide variability between transcriptome coverage in individual cells, with a small number of cells covered very highly (>5000 genes detected) and the majority covered at a shallow level (<1000 genes detected; Fig. 1E). This distribution of transcriptome coverage was markedly more uniform for the Fluidigm data, suggesting that the coverage variability in DropSeq data results from experimental factors such as variability between droplets or beads.

It seems likely that per-cell transcriptome coverage will affect estimates of population statistics such as mean or variability measures, or the ability to assign cells to specific cell states, but it is unclear how much transcriptome coverage is needed for these applications. This raises the question of how stringently to filter cells in order to obtain sufficient quality for particular biological inferences, which is difficult to do without a ground truth reference. We therefore set about answering this question by analyzing subsets of the DropSeq data with different per-cell transcriptome coverage levels, and using RNA FISH as an independent gold standard.

First, we compared the mean level of RNA detected across all cells by comparing unique molecular identifiers per million (in the case of DropSeq) or reads per million (in the case of Fluidigm) to average counts per cell from RNA FISH (Fig. 1F). Given that the mean is less dependent on single cell variability, we expected to find a general correspondence between RNA FISH and the RNA sequencing-based methods. We found that the correlation was fairly strong for both methods (DropSeq R = 0.61; Fluidigm R = 0.63), and this correlation was similar across a range of sequencing depths (Supp. Fig. 1D).

Interestingly, we noted that despite the overall correspondence in mean transcript abundance between RNA FISH and RNA sequencing (see also [18,19]), the differences between the single cell RNA sequencing methods and RNA FISH were shared between both comparisons, and that these discrepancies were of similar magnitude and direction. This raises the question whether these expression differences result from differences in sample handling or from methodological differences between RNA FISH and RNA sequencing. We compared the expression of these genes between DropSeq and Fluidigm (Fig. 1G), and found a much stronger correlation (R = 0.95). Furthermore, the correlation between these methods and bulk RNA sequencing (also reads per million) was similarly high (DropSeq R = 0.94; Fluidigm R = 0.92), even though the bulk RNA sequencing was performed on RNA harvested directly from adherent cells as opposed to the dissociated cells used for DropSeq and Fluidigm (Fig. 1F). Given the radical differences in RNA isolation and library preparation between these three methods of RNA sequencing, we concluded that the differences between RNA FISH and RNA sequencing transcript level estimation likely stem from systematic biases in sequencing itself and not from biases introduced by the different protocols or sample handling.

We next sought to compare the single cell distributions of transcript abundance as measured by either single cell RNA sequencing and RNA FISH. We first used the Kolmogorov–Smirnov (KS) statistic, which measures the deviation between two expression frequency distributions. We calculated the KS statistic between the single cell datasets and RNA FISH for each gene (Supp. Fig. 2A). We found that the KS statistic improved slightly and continuously as the transcriptome coverage threshold increased, but there was no clear point demarcating a value above which the results were qualitatively more consistent (Supp. Fig. 2B).

While KS provides a metric to compare distributions in general, it may not as effectively capture differences in specific features of distributions that may be of interest; thus, other metrics may provide a more sensitive point of comparison in certain contexts. Because we were specifically interested in detecting rare-cell expression patterns, we turned to the Gini coefficient, developed by Corrado Gini as a means of quantifying income inequality. In the context of single cell expression level [20], a Gini coefficient of zero signifies an equal distribution where all cells express exactly the same levels of a particular gene, whereas a Gini coefficient of one signifies the most extreme level of jackpot expression in which all the RNA is concentrated in a single cell while all the others have none. Intermediate Gini coefficients correspond to intermediate levels of rare-cell heterogeneity (Fig. 2A).

**Figure 2.**
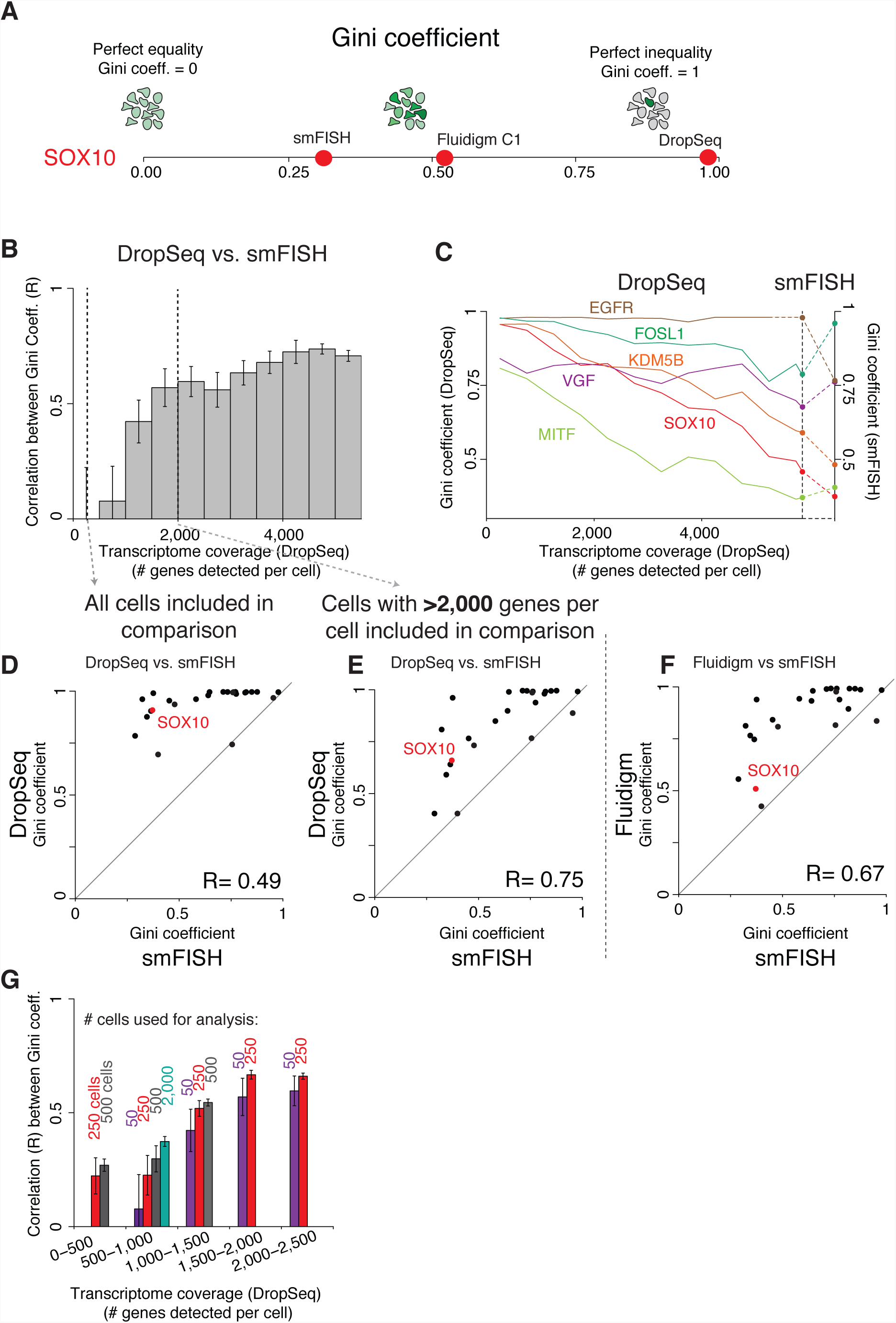
Measurement of gene expression heterogeneity in scRNA-seq is highly dependent on transcriptome coverage. (A) The Gini coefficient measures a gene’s expression distribution and captures rare cell population heterogeneity. The Gini coefficient is often overestimated by scRNA. (B) Correlation between Gini coefficients measured through DropSeq and smFISH across different levels of transcriptome coverage (# genes detected per cell). Error bars represent ± 1 standard deviation across bootstrap replicates. (C) Gini coefficient for five genes measured by DropSeq (left y-axis) at different levels of transcriptome coverage as well as by smFISH (right y-axis). (D,E) Scatterplot of the correspondence between Gini coefficients for 26 genes measured by both DropSeq and smFISH. The correlation between estimates improves when only cells with high transcriptome coverage (>2,000 genes per cell) are included in the analysis. (F) Scatterplot of the correspondence between Gini coefficients for 26 genes measured by Fluidigm and smFISH. (G) Correlation between Gini coefficient estimates measured by DopSeq and smFISH using different population sizes (# of cells) and levels of transcriptome coverage. Error bars represent ± 1 standard deviation across bootstrap replicates.

The genes whose expression we analyzed by RNA FISH had expression Gini coefficients ranging from 0.29 to 0.98, with housekeeping genes such as *GAPDH* having a Gini coefficient of 0.33 while resistance markers like *EGFR* and *WNT5A* had high Gini coefficients of 0.76 and 0.83. We then wondered how accurate single cell RNA sequencing measurements of Gini coefficients would be, given the technical sensitivity of single cell RNA sequencing. Indeed, we found that the Gini coefficients from both platforms were generally far higher than in RNA FISH; for instance, *SOX10* has a Gini coefficient of 0.38 by RNA FISH, but has a Gini coefficient of 0.91 by DropSeq and 0.51 by Fluidigm (Fig. 2A).

Given that the Gini coefficients calculated from Fluidigm (which had higher mean library complexity as defined by coverage per cell) corresponded more closely to Gini coefficients from RNA FISH than did DropSeq (Fig. 2D,F,C), we reasoned that accuracy of Gini estimation may depend on transcriptome coverage. Single cell RNA sequencing is often plagued by so-called zero inflation, in which some cells artificially yield low or zero levels of many transcripts [10,21,22], which would likely inflate estimates of Gini. To test this hypothesis, for each of the genes for which we had RNA FISH data, we computed the Gini coefficients for Drop-seq samples binned by the number of genes detected per cell. We found that the Gini coefficient estimates for genes with low variability (e.g. SOX10) generally decreased as the number of genes detected per cell increased, while the Gini estimates for highly variable genes (e.g. EGFR) remained high (Fig. 2C). Overall, as the stringency threshold for transcriptome coverage increased, the correspondence with the Gini coefficients measured by RNA FISH increased (Fig. 2B), with the highest quality cells giving a similar correlation for both Gini coefficient and mean expression. Upon calculating the correlation coefficient between Gini coefficients as measured by single cell RNA sequencing vs. RNA FISH over a range of number of genes detected per cell, we found that the correspondence for DropSeq increased most until the number of genes detected per cell reached around 2000, but did not increase appreciably beyond that number (Fig. 2B). This suggests that a threshold for transcriptome coverage of 2000 genes detected per cell (for our DropSeq data) was sufficient for reasonably accurate quantification of rare-cell expression via single cell RNA sequencing. (Notably, cells in the Fluidigm dataset generally had more uniform (and uniformly higher) number of genes detected (Fig. 1E), and as expected, the Gini coefficients calculated from these data were more accurate (Fig. 2F).)

The improvement of Gini coefficient estimates from stringently filtering the DropSeq data are driven by decreases in artificially high Gini coefficients; i.e., the lowering of Gini coefficients for genes that are relatively uniformly expressed as the transcriptome coverage increases. This leads to the somewhat counterintuitive prediction that having a small number of cells with higher transcriptome coverage leads to more accurate Gini coefficients for rare-cell expression than a large number of cells with shallow transcriptome coverage, because doing so reduces the false positive high Gini coefficient genes. We tested this by estimating the Gini coefficient for a range of sample sizes (number of cells) for cells binned by number of genes detected per cell (Fig. 2G). We found that increased sample size did improve the similarity of our Gini estimate with smFISH estimate. We also found that using a large number of cells with low transcriptome coverage (eg. n = 2000 with 500-1,000 genes detected) still provided a worse estimate than using a small number of higher complexity cells (eg. n=50 cells with 1,500-2,000 genes detected). This suggests that having a pool of few high-transcriptome coverage cells may yield more accurate Gini coefficients due to reduction of artificially high Gini coefficients that stem from the low transcriptome coverage of shallow depth single cell RNA sequencing.

Given this guidance for determining a threshold for whether to include a cell from the DropSeq data based on RNA FISH, we then proceeded to ask whether such a threshold on transcriptome coverage would be useful for cell-state determination, and whether such a threshold could be identified in a context in which RNA FISH data is not available, focusing on the cell cycle as a canonical example [17]. Our analysis consisted of classifying cells based on their expression of a panel of genes known to be associated with cell cycle [23]. We first classified our DropSeq cells for cell cycle phase (Fig. 3B, top). To assess our ability to detect biological signal, we also generated a null expectation, classifying cell cycle phase for randomly permuted data (Fig. 3B, bottom) (data permuted by, independently for every gene, randomly assigning transcript counts between all cells). We found no significant difference between the two. To assess the relationship between transcriptome coverage and our ability to detect biological signal, we classified cells binned by number of genes detected per cell, and then we increased the stringency threshold and measured how signal emerged above the randomized control. We defined signal strength as the difference between how well a cell’s transcriptome correlated with the idealized signature of the assigned phrase and how well it correlated with the “opposite” phases (e.g., for a G1/S-assigned cell, how well it correlated with G1/S minus how well it correlated with G2 and G2/M phases) (Fig. 3D). We found that our ability to detect cell cycle phase improved with the number of genes detected per cell. Moreover, we again found that at a threshold of around 2000 genes detected, the signal strength significantly increased above randomized control (Fig. 3E), and the number of cells in each phase of the cell cycle reached more plausible values (Fig. 3A). The classification of cells differed much more from randomized data when cells were thresholded above this stringency level (Fig. 3C), and fit more with the canonical view of the cell cycle, with most cells in G1 phase and a smaller number in G2 or M. An important consideration, however, is that the ability to classify the biological state of a cell likely depends on the number genes that mark a given state. Are higher transcriptome coverages required for more subtle biological states? To test this, we classified cell cycle phase using random subsets of phase marker genes, for a range of transcriptome coverages (genes detected per cell) (Fig. 3F). As expected, our ability to detect biological signal increased with the number of marker genes available, while for more subtle biological signals, higher transcriptome coverages were required to distinguish the signal from random. For example, distinguishing a cell cycle phase based on the expression of 30 marker genes requires a transcriptome coverage of >1500 genes/cell, while a cell state defined by 10 marker genes requires >4000 genes/cell for the detection of any reliable signal (Fig. 3F).

**Figure 3.**
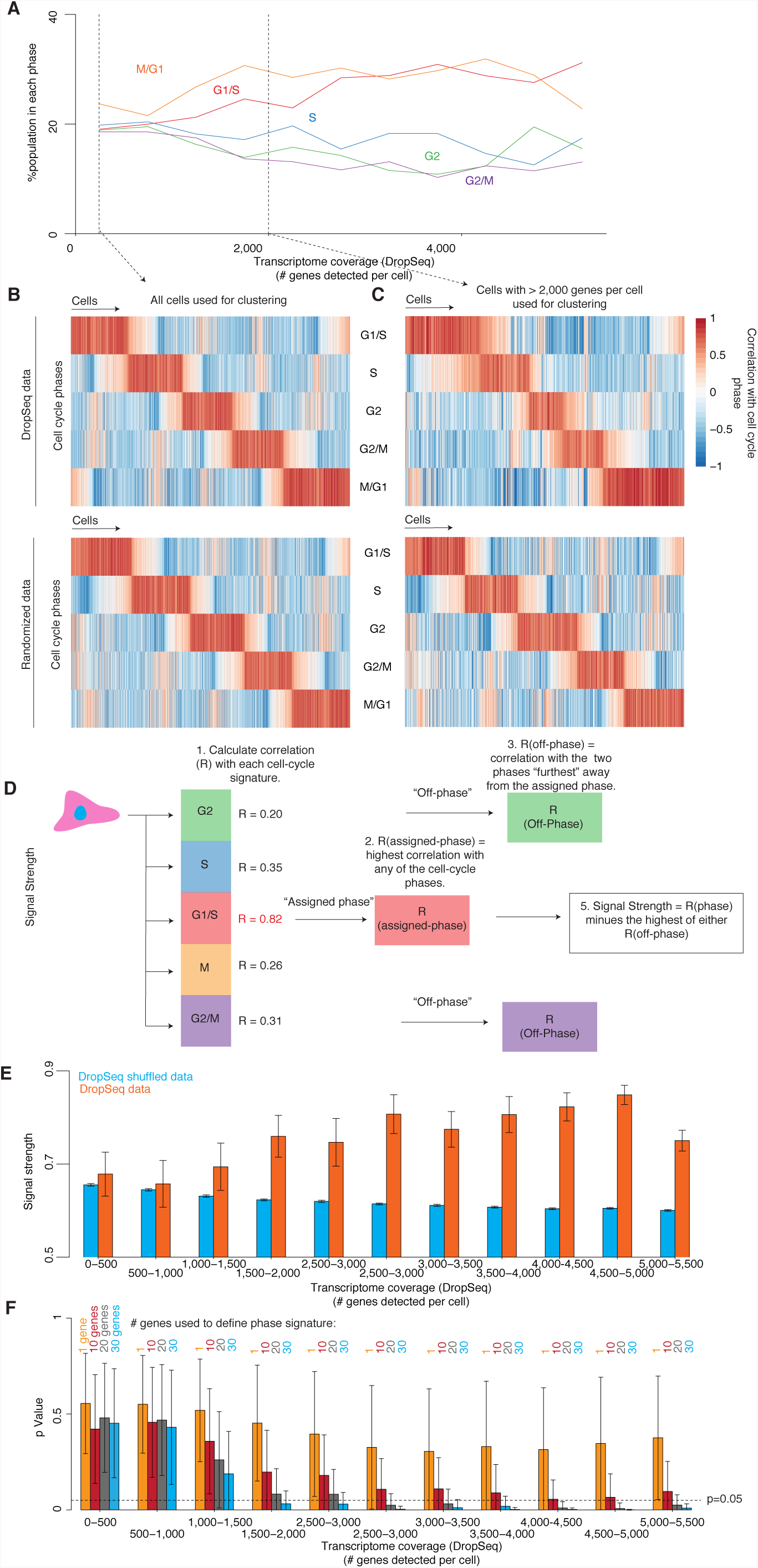
Correct classification of single cells into multi-genic states is highly dependent on transcriptome coverage. (A) Percent of cells assigned to a cell cycle phase (M/G1, G1/S, S, G2, or G2/M) at different levels of transcriptome coverage (# genes detected per cell). (B, C) Correlation of a cell’s gene expression signature (columns) with each of the cell cycle phases (rows) for the DropSeq dataset (top) as well as for a null model (bottom) where the expression level of all genes in a given cell were randomly shuffled across cells. We did this analysis including either all cells (B) or only cells with a high transcriptome coverage (C). (D) Signal Strength allows us to measure how strongly and uniquely a cell correlates with a given cell cycle phase. (E) Signal strength across different levels of transcriptome coverage for DropSeq (orange) and a null model of randomized DropSeq data (orange). Error bars represent ± 1 standard deviation across bootstrap replicates. (F) p-value of signal strength at different levels of transcriptome coverage using a different number of genes to characterize the phase. Bar height indicate mean across bootstrap replicates. Error bars represent ± 1 standard deviation across bootstrap replicates.

These results show that biological inferences of cell state—just as with rare-cell analysis—also require cells of sufficient transcriptome coverage, with a similar threshold level for stringency.

## Discussion

Single cell RNA sequencing is a transformational tool for studying gene expression in single cells, but as yet, little guidance exists on what criteria on such datasets are required for making reliable inferences for different biological questions. Partly, this is because of a lack of complementary gold-standard data. Here, we have used large-scale single molecule RNA FISH datasets as a gold-standard, focusing in particular on the question of rare-cell variability in gene expression. We found that DropSeq data produced cells with a wide variety of transcriptome coverages, and that effective identification of rare-cell expression required limiting analysis to those cells with higher transcriptome coverage. Using these same quality thresholds, we were also able to more robustly classify cells by position in cell cycle.

This work highlights the tradeoffs inherent to the analysis of single cell RNA sequencing data. With DropSeq (but also for other methods), the question arises as to how deep the transcriptome coverage for any one cell must be before including it in the analysis, because the vast majority of putative cells have very shallow transcriptome coverage, either due to amount of sequencing performed or inherent library complexity. In the context of the literature, our results suggest that these choices strongly depend on the biological question under consideration. Many applications of single cell RNA sequencing have been for the identification of different cell types in tissues, and for such questions, a shallow transcriptome coverage may be sufficient, essentially because cell types typically differ in the expression of hundreds or thousands of covarying genes, any subset of which are sufficient for classification [11]. Our results show that distinguishing more closely related cell types or cell states which differ by ∼1-30 genes, such as the position within cell cycle, requires higher transcriptome coverage for accurate determination. Similarly, we found that even moderately accurate classification of a particular gene as expressing in a rare-cell manner (i.e., high Gini coefficient) requires considerably deeper transcriptome coverage per cell. In general, our results demonstrate that an accurate statement about the single cell variability of any particular gene or small subset of genes requires higher transcriptome coverage than that needed for cell-type identification studies.

How might one determine such a threshold? In our case, we were able to use single molecule RNA FISH data to provide a ground truth as a basis for comparison, but such data is typically not available to cross-reference against, and it is likely that the threshold may vary between sample types and even cell lines. We suggest a procedure in which one repeats the analysis while varying the stringency of the threshold for including cells in the analysis and then measuring the strength of the signal for the comparisons of interest (such as how distinct cell populations are compared to randomized data). If the cells are actively proliferating, then one could perform an analysis similar to that performed in Fig. 3 for cell cycle to help calibrate the threshold, while realizing that the ideal threshold may not be directly transferable because it will depend on factors such as the number of differentially expressed genes and magnitude of expression differences for each gene.

In this vein, there are also new computational tools such as MAGIC [24] that aim to recover correlations from shallow-coverage cells in single cell RNA sequencing datasets, as well as tools like SAVER [25] (co-submitted manuscript) that are even able to recover distributions from shallow-coverage cells in single cell RNA sequencing datasets. SAVER is able to recover these distributions by training a prediction model across all cells regardless of coverage, thus yielding counts per cell based on a weighted average of the model prediction and the experimental observations. It remains to be seen how much these methods rely on the particulars of the distribution of transcriptome coverage across cells. It is also interesting to consider the possibility of both sequencing a few cells at very high coverage and a much larger number of cells at a shallower read depth to see if it is possible to recover more information from these combined datasets.

## Acknowledgements

We thank Emily Shields for performing some preliminary analysis, and members of the Murray and Raj lab for helpful comments. JM and HD acknowledge support from R21 HD085201. AR and ET acknowledge support from NIH New Innovator Award DP2 OD008514, NIH/NCI PSOC award number U54 CA193417, NSF CAREER 1350601, NIH R33 EB019767, P30 CA016520, NIH 4DN U01 HL129998, NIH Center for Photogenomics RM1 HG007743, and the Charles E. Kauffman Foundation (KA2016-85223). SMS acknowledges NIH F30 AI114475. R.B. acknowledges support from the NIH (DP2MH107055), the Searle Scholars Program (15-SSP-102), the March of Dimes Foundation (1-FY-15-344), a Linda Pechenik Montague Investigator Award, and the Charles E. Kauffman Foundation (KA2016-85223).

## Methods

### Experimental methods

Cell culture - We obtained WM989 melanoma cells from the lab of Meenhard Herlyn and derived A6 and A6-G3 subclones in our lab. We grew them in Tu2% media (78% MCDB, 20% Leibovitz’s L-15 media, 2% FBS, and 1.68mM CaCl2). We grew 3T3 murine cells in DMEM media (10% FBS, 0.5% pen/strep), and murine JC4 suspension cells in IMDM media (10% FBS, 2% pen/strep, 50ng/ml Kit ligand, 2 U/ml erythropoietin, 4.5 × 10^-5 M monothioglycerol).

RNA FISH - For RNA FISH we seeded cells in two-well LabTek chambered coverglasses and cultured them to ∼50-70% confluency. We performed single molecule RNA FISH and high-throughput microscopy scans as previously described [7] (and Shaffer et al. Nature in press). In short, we first fixed adherent cells with 4% formaldehyde in PBS for 10 min at RT and permeabilized with 70% EtOH at 4C overnight. We hybridized FISH probes (DNA oligonucleotides conjugated to fluorescent dyes) overnight at 37C, washed away unbound probes, and stained DNA with DAPI prior to acquiring a tiled grid of images. Note that in our imaging system, we measured expression in a single z-plane of the cell; thus, the exact numbers for each cell are not total mRNA counts per cell, but rather an amount proportional to the total. For iterative FISH, we stripped DNA probes and hybridized new ones as in Shaffer et. al. (Nature, in press) In short, after an initial round of imaging, we removed bound DNA probes using 60% formamide on 2X SSC during a 15 minutes incubation at 37C. We then removed the formamide with three 15-minute washes at 37. Finally, we washed one last time with wash buffer prior to adding a new set of DNA probes.

Dropseq - We generated single cell suspensions by trypsinizing adherent cells with 0.05% trypsin-EDTA or by harvesting suspensions cells. We passed all cells through a 40 micron filter and diluted them to 100 cells/ul in PBS-BSA. We carried out all subsequent steps as detailed by Macoscko et. al, protocol v3.1 (http://mccarrolllab.com/dropseq/). In short, we loaded cells in PBS-BSA and barcoded beads (chemgenes Barcoded Bead SeqB, cat. No. MACOSKO-2011-10) in lysis buffer onto a droplet generating microfluidic device. After breaking the droplets, we pooled the beads into aliquots of ∼60,000, reverse transcribed the RNA captured by the barcoded beads, and digested unbound poly-dT tails via exonuclease treatment. We PCR-amplified STAMPs (2000 beads per reaction), purified cDNA using AMPure beads and quantified the library via Agilent’s High Sensitivity DNA Chip. We then tagmented the resulting cDNA with nextera XT adapters and purified the final library with Ampure beads. We sequenced all libraries using Nextseq 500 with a custom Dropseq read 1 primer.

Fluidigm - To prepare single cell suspensions, we dissociated WM989-A6-G3 cells as above. We immunostained the cells as per Shaffer et. al (Nature, in press). Briefly, we incubated cells for 1 hour at 4C with 1:200 mouse anti-EGFR antibody, clone 225 (Millipore, MABF120) in 0.1% BSA PBS. We then washed twice with 0.1% BSA-PBS and then incubated for 30 minutes at 4C with 1:500 donkey anti-mouse IgG-Alexa Fluor488 (Jackson Laboratories, 715-545-150). We washed the cells again with 0.1% BSA-PBA and incubated for 10 minutes with 1:500 anti-NGFR APC-labelled clone ME20.4 (Biolegend, 345107). After we washed the cells with 0.1% BSA-PBS and pelleted them, we resuspended them in Tu2%, passed them through a 35 micron filter, and diluted them to a final concentration of ∼350 cells per ul in Tu2%. We prepared the samples and sequencing library according to the manufacturer’s instructions (https://www.fluidigm.com/products/c1-system). In short, we loaded and captured single cells on Fluidigm’s C1 integrated fluidic circuit and inspected the capture chambers via microscopy. We then lysed the cells, barcoded the captured mRNA via RT with a barcoded primer, and amplified the resulting cDNA via PCR. Unlike DropSeq, this protocol uses no unique molecular identifiers to label RNA molecules. After we harvested the amplified cDNA, we tagmented the library using Nextera’s XT DNA sample preparation kit (following Fluidigm’s version of the protocol), purified the final library using Ampure beads and quantified using Agilent’s High Sensitivity DNA Chip. We sequenced the library using a Nextseq 500.

### Analysis methods

#### Drop-seq alignment and quantification

Initial Drop-seq data processing was performed using Drop-seq_tools-1.0.1 (http://mccarrolllab.com/dropseq/), and following protocol described in seqAlignmentCookbook_v1.1Aug2015.pdf, accessed from the same site. Data were aligned using STAR version 2.4.2a, downloaded from github on Jan 21, 2016. Data were aligned to reference genome builds hg38 (Human) and mm10 (Mouse), and using reference transcriptome annotations Gencode21 (Human) and Refseq mm10 (Mouse), concatenated with ERCC sequences. Reference transcriptome annotations Gencode21 (Human) and Ensembl mm10 release 83 (Mouse), concatenated with ERCC annotations. Briefly, reads with low-quality base in either cell or molecular barcode were filtered and reads were trimmed for contaminating primer or poly-A sequence. Sequencing errors in barcodes were inferred and corrected, as implemented by Drop-seq_tools-1.0.1. Uniquely mapped reads, with <= 1 insertion or deletion, were used in quantification. To account for differences in molecule recovery, cell measurements were normalized to UMI per million (UPM).

#### Fluidigm alignment and quantification

Fluidigm sequence data were demultiplexed using mRNASeqHT_demultiplex.pl (https://www.fluidigm.com/c1openapp/scripthub/script/2015-08/mrna-seq-ht-1440105180550-2). Demultiplexed data were processed in the same manner as Drop-seq data, with a few modifications: 5’ ends of reads were not trimmed, and reads (rather than UMI) were used for quantification. To account for differences in molecule recovery and sequencing depth, cell measurements were normalized to reads per million (RPM).

#### Bulk sequencing alignment and quantification

We sequenced mRNA in bulk from WM989-A6 populations as per Shaffer et. al. We isolated mRNA and built sequencing libraries using the NEBNext Poly(A) mRNA Magnetic Isolation Module and NEBNext Ultra RNA Library Prep Kit for Illumina. We sequenced the libraries either on a HiSeq 2000 or a NextSeq 500 to a depth of approximately 20 million reads. We then aligned the reads to hg19 and quantified reads per gene using STAR and HTSeq.

#### FISH quantification

All image analysis was performed as per Shaffer et. al. We developed a MATLAB analysis pipeline that segments nuclei of individual cells using DAPI images. The pipeline then identifies regional maxima as potential DNA FISH spots and assigns them to the nearest nuclei. We then select a signal intensity threshold for each RNA FISH channel to differentiate background from RNA FISH signal. We then extract the position of every cell in the scan and the number of RNA molecules for each fluorescent channel. To match cells across subsequent hybridizations, we developed a software that shifts cells in the first hybridization to all potential candidates in the subsequent hybridization. It then chooses the best match as the one that minimizes the total distance for nearby cells. We then matched cells by proximity and discarded those cells that did not match uniquely to a nearby cell.

#### Selecting quality single cell Drop-seq data

Cell barcodes were classified as quality human cells, based on the following criteria: 1) Greater than 80% of species-specific transcripts were assigned to human, and at least 100 species-specific transcripts were available for assignment. 2) The cell barcode was not assigned a synthesis error. The remaining barcodes were filtered to retain the expected number of cells. Based on experiment, we expected 8640 single human cells, and we retained the 8640 cell barcodes with largest read depth (Supplemental Table 1).

#### Selecting quality single cell Fluidigm data

The Fluidigm system allowed cells to be imaged before processing, and for images to be associated with sequencing data. In our automated setup, not all wells were imaged, and well numbers were not captured in images (though column identity on the chip was known). In order to use images to identify quality single cells, we first re-ordered visual annotations to best match read depth observed per well (so that images of empty wells had low depth compared to images with individual or multiple cells). Given re-ordered images, wells were classified as quality single cells if: 1) based on associated image, the well was annotated as containing a good, single cell, and 2) if the cell appeared to be distinct from wells annotated as empty by read depth. Both criteria were required (Supplemental Tables 2 and 3).

#### Sufficiency of sequencing depth

We wanted to ensure that our metric of library complexity (number of genes observed) did not reflect sequencing depth. To test whether the Drop-Seq and Fluidigm experiments were sequenced to a sufficient depth, we examined the relationship between experimental read depth and the average number of genes observed in single cells. To do this, we randomly and uniformly subsampled reads from the Drop-seq (or Fluidigm) read counts table, for a variety of experimental sequencing depths. We generated 10 random samples at each sequencing depth, and report the average number of observed genes, across cells and sample replicates. We generated random samples for an average depth per cell of 100, 500, and 1000–500,000 (step size of 1000) raw reads per cell. (For Drop-seq data, our experimental depth allowed testing depths up to 120,000 average raw reads per cell.) To identify the number of reads to subsample from the read count table, given these raw read depths per cell, we calculated the fraction of all sequenced reads that were assigned to the read count table. At each selected experimental depth, we used this fraction of reads to subsample the read counts table. For Dropseq data, 11.2% of raw reads were uniquely assigned to genes in quality cells. In Fluidigm data, 29.3% of reads were uniquely assigned to genes in quality cells.

#### Tissue-marker gene expression

We selected tissue marker genes for melanocytes, pancreas, heart, and spleen from TIGER (http://bioinfo.wilmer.jhu.edu/tiger/). For each tissue type the genes were selected for analysis based on the expression level in their respective tissue and their presence in both single cell RNA sequencing datasets.

#### Comparison of average measurements

We calculated mean ± 2 SEM for each measurement type. Two genes with FISH measurements were excluded (VGF and NGFR) due to unreliable FISH measurements. One additional gene (AXL) was not observed in the Fluidigm data, and was excluded from Fluidigm comparison. Pearson correlations were calculated over genes observed in both Drop-seq and Fluidigm and were calculated on a log_10_ scale.

#### Filtering and normalization for calculation of rare cell variability

It has been shown that a portion of molecular variability across single cells is due to cell volume [18]. To focus on rare cell variability, we normalized smFISH cells to GAPDH, using GAPDH levels as a proxy for cell volume. Cells with <50 GAPDH molecules observed were filtered prior to normalization. For visualization, we scaled normalized values by 400 so that normalized counts were on roughly the same scale as single cell molecular counts. In order that sequencing data remain comparable to smFISH data, we filtered Drop-seq and Fluidigm cells with no observed GAPDH. We then scaled the (sequencing-depth normalized) Drop-seq and Fluidigm data so that the median GAPDH level across cells was 400, so that sequencing measurements were on a similar scale to smFISH measurements.

#### Measure of rare cell variability

Gini coefficients were calculated using the R package “ineq”.

#### Effect of library complexity on Gini coefficient estimate

To test the effect of library complexity on estimates of population statistics, we binned cells by library complexity, using the number of observed genes as our metric of complexity. We used bins ranging from 0 to 5500 observed genes, with a step size of 500 genes. Sample size (the number of cells in a bin) is expected to affect the estimate of the Gini coefficient. We controlled for sample size (number of cells) by randomly subsampling cells within a bin, to reach 50 cells per bin, prior to calculating the Gini coefficient. Random sampling was repeated 100 times per bin. So that normalization was consistent across all random subsamples, we normalized all cells to cellular GAPDH level. As previously, smFISH cells with <50 GAPDH molecules were excluded, as were Drop-seq and Fluidigm cells with no GAPDH observed. For each complexity bin, we calculated the Pearson correlation of Gini coefficient estimates calculated on Drop-seq data with those calculated using smFISH data. For each bin, we report the average correlation across subsample replicates ± 1 standard deviation.

#### Effect of sample size (number of cells) on Gini coefficient estimate

To evaluate the effect of sample size on Gini coefficient estimates, we repeated the analysis described above for a variety of numbers of cells for each complexity bin.

#### Cell cycle phase classification

To assess our ability to detect biological expression patterns using Drop-seq and Fluidigm measurements, we assigned cell cycle phase to individual cells, following the approach used in Macosko et al. [17] and using cell cycle marker genes identified in Whitfield et al. [23]. The Macosko et al. approach involves the following steps: 1) Marker genes were filtered to exclude genes that do not cycle in melanoma cells. For each set of genes assigned to a particular cell cycle phase, the average expression profile was calculated across the data set. For each individual gene within that set, the correlation with this average profile was calculated. Genes with correlation <0.3, in either Drop-seq or Fluidigm data, were excluded. 2) Depth-normalized read counts were zero-adjusted and log_2_ normalized. 3) For each cell and phase, a phase score was assigned by calculating the average normalized value across marker genes for that phase. 4) Phase scores were z-normalized, first across cells within each phase, and then across phases within each cell. 5) Sample phase was assigned to each cell. To do this, a binary score profile was created for idealized cells at each phase and phase transition. The correlation of a cell’s normalized score profile with this set of idealized profiles was calculated. A cell was assigned a phase based on the maximum observed correlation.

#### Effect of library complexity on cell cycle phase classification

To test the effect of library complexity on cell phase classification, we binned cells by the number of observed genes as described above and, as above, we controlled for sample size (number of cells) by randomly subsampling cells within a bin, to reach 50 cells per bin. For each set of sampled cells, we classified cell cycle phase as described above. To generate a random expectation for cell cycle phase categorization, we generated 1000 random counts tables, shuffling counts across cell cycle marker genes for each sample. For each table, we proceeded with cell cycle phase classification as described above. We used the same sets of samples as used in test data, so that each tested population of cells was compared to a biologically and technically matched population with randomized expression profiles.

To summarize the strength of biological signal, we calculated for each cell the best correlation with an idealized phase profile (the assigned phase for the cell) and the best correlation with an idealized phase profile for an “off” phase, or a cell cycle phase that does not neighbor the assigned phase. We report the difference between these correlations. To provide a population-level statistic, we calculate the average strength of biological system for each tested population (each randomly sampled set of 50 cells). To assess the significance of this statistic, we calculated the same statistic for each null (randomly shuffled) population, and report the fraction of times that a signal as large or larger is observed. Finally, we summarize these results across the randomly sampled populations (the random sets of 50 cells), reporting the average ± 1 standard deviation across subsample replicates.

#### Effect of number of marker genes on cell cycle phase classification

To evaluate the effect of the number of available marker genes, we repeated the analysis described above for a variety of numbers of genes. We randomly selected *n* marker genes for each phase (*n* from 1 to 30), and then ran the analysis described above on that subset of genes. We repeated this 100 times for each *n*. For randomized data, we selected *n* genes for each phase once, because the identity of the gene has been lost in randomizing the data. (This means each of the 1000 randomized replicates is essentially a different random gene set sample as well.) So, each of the 100 replicates of test data for a given *n* are compared to the same null expectation.

---

**Supplemental Figure 1.**
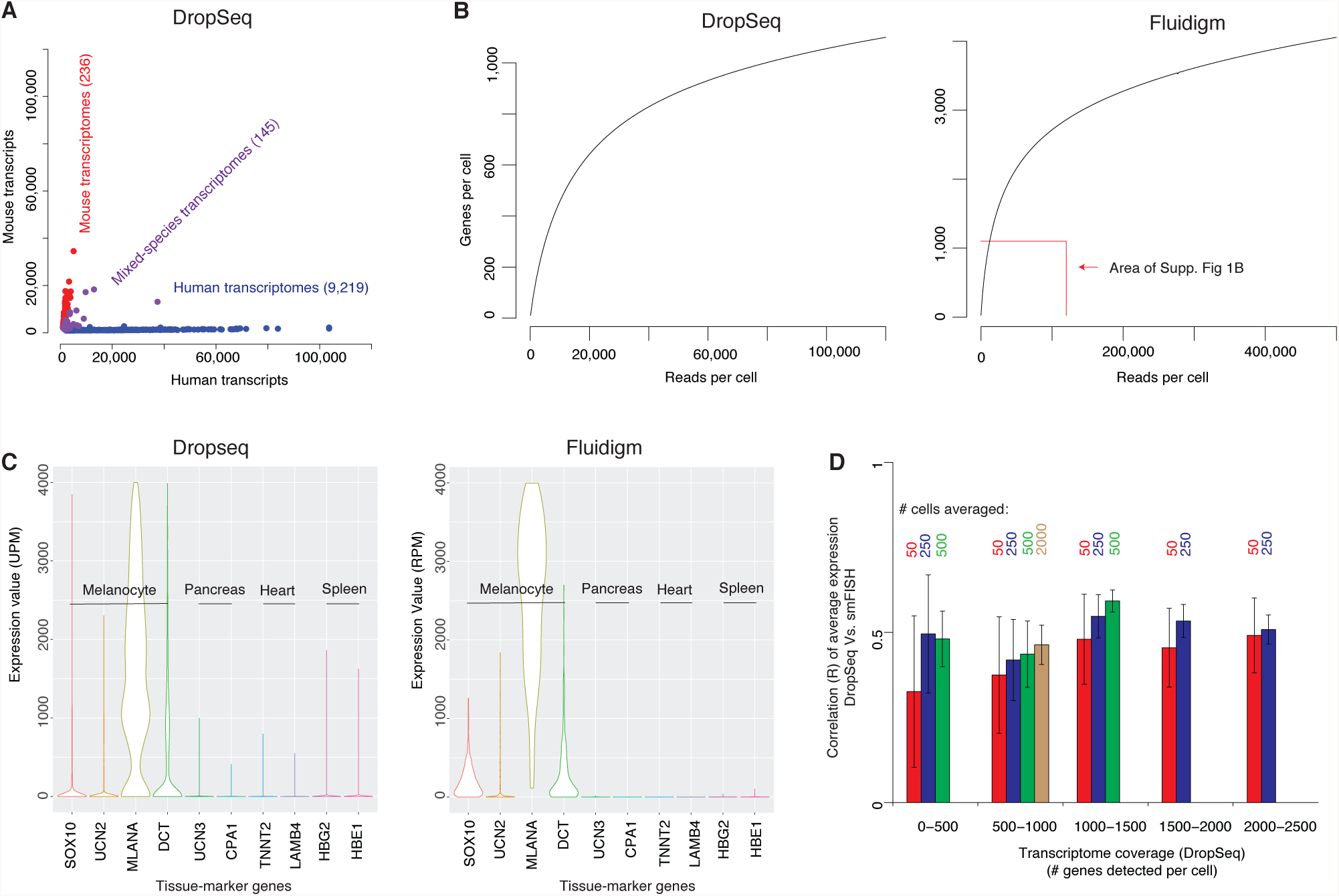
(A) Scatterplot representing the number of human and mouse transcripts associated with each cell barcode in the DropSeq dataset. (B) Sequencing coverage measured by the number of genes detected per cell as a function of the number of reads obtained for each cell for DropSeq (left) and Fluidigm (right). (C) Gene expression estimates of tissue-marker genes for DropSeq (left) and Fluidigm (right). (D) Correlation of average gene expression estimates between DropSeq and smFISH at different levels of transcriptome coverage (# genes detected per cell) using four different population sizes (50, 250, 500, and 2000). Not all population sizes are available at all levels of transcriptome coverage. Error bars represent ± 1 standard deviation across bootstrap replicates.

**Supplemental Figure 2.**
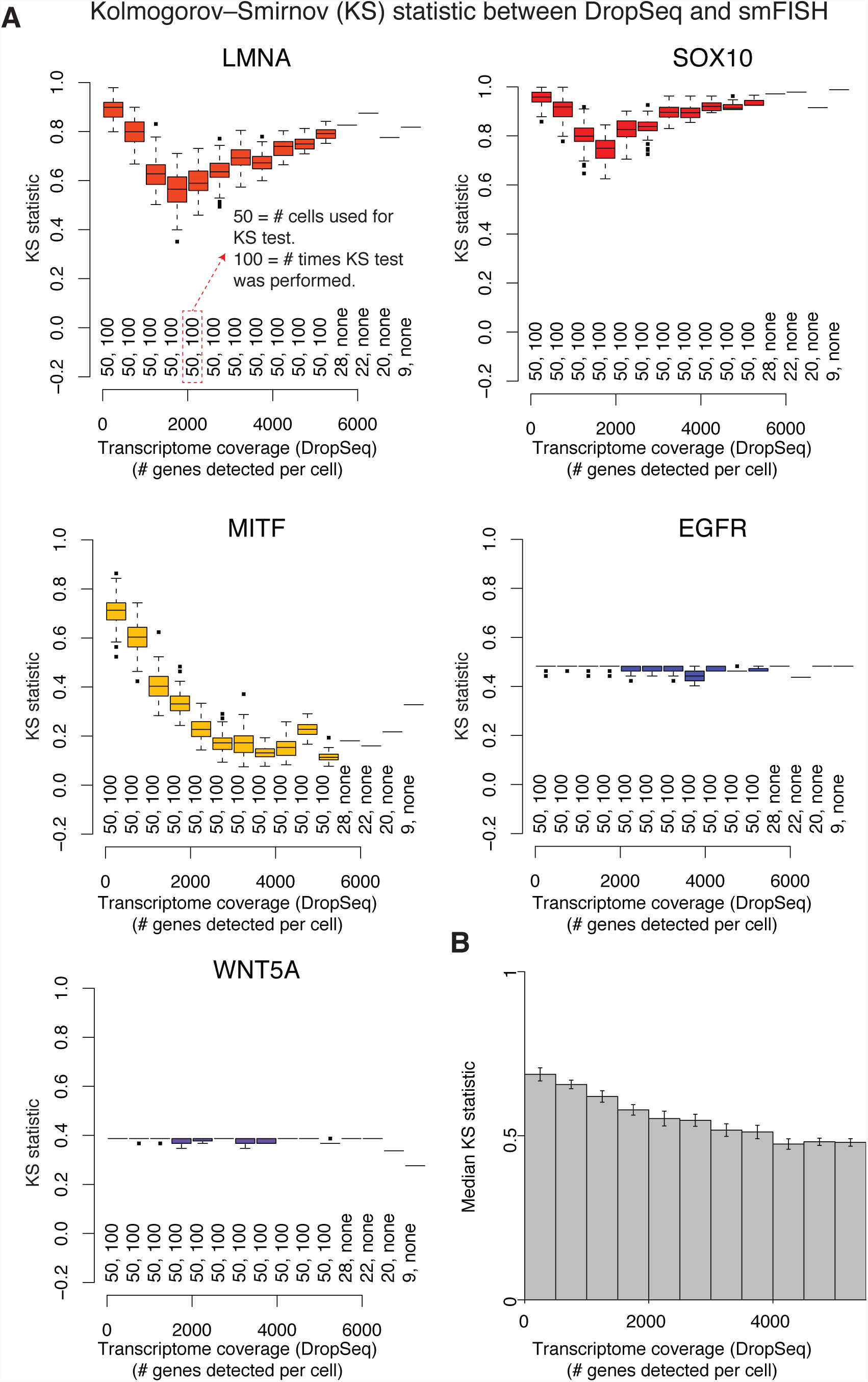
(A) Comparison of the gene expression distribution (Kolmogorov-Smirnov statistic) for five genes (LMNA, SOX10, MITF, EGFR, and WNT5A) measured by DropSeq and smFISH. We measured the KS statistic across different levels of transcriptome coverage (# genes detected per cell). Unless otherwise indicated, at each level of transcriptome coverage the KS test was repeated 100 times, each time randomly sampling 50 cells from the population. The distribution of KS values across bootstrap replicates is depicted as a boxplot. (B) Median KS statistic between DropSeq and smFISH of the 26 genes listed in Fig. 1A across varying degrees of transcriptome coverage. Bar height indicates the average across bootstrap replicates. Error bars represent ± 1 standard deviation across bootstrap replicates.

